# Anticipatory Threat of Shock Reduces the Number of Items Reported From a Brief Visual Display: Evidence for Capacity Loss Along the Scanning Order

**DOI:** 10.64898/2026.11.01.763403

**Authors:** Barış Akar, Merve Aydın Karabaş, Tuğba Altıntaş

**Affiliations:** Department of Neuroscience, Institute of Health Sciences, Üsküdar University, İstanbul, Türkiye; Department of Perfusion, Anaesthesia Program, Vocational School of Health Services, Üsküdar University, İstanbul, Türkiye; Department of Health Management, Faculty of Health Sciences, Üsküdar University, İstanbul, Türkiye

**Keywords:** threat of shock, anticipatory anxiety, iconic memory, whole report, visual short-term memory, attentional narrowing

## Abstract

**Background:** Anticipatory anxiety impairs working memory, but whether it also limits the earliest, high-capacity stage of visual storage is unclear. We asked whether the mere expectation of an aversive electrical stimulus reduces how many items people can report from a briefly flashed display, and whether any loss is spatially uniform.

**Methods:** Eighty-eight healthy adults aged 18–25 years were randomly allocated to a threat-of-shock condition (surface electrodes on the wrist and the instruction that a painful stimulus might be delivered; none was) or a control condition (*n* = 44 each). Participants viewed a 4 × 3 letter matrix for 400 ms and performed untimed written whole report. Correct recall (0–12), intrusions and the position of each reported letter were scored. Groups were compared with Welch’s *t* test, the Mann–Whitney *U* test, bootstrapped effect sizes, a binomial generalised linear model and a Bayes factor; spatial effects were tested with generalised estimating equations (GEE) accounting for letters nested within participants.

**Results:** The threat group reported fewer correct letters (3.41 ± 1.09) than controls (4.70 ± 1.29); Welch *t*(83.6) = 5.11, *p* < .001, Hedges’ *g* = 1.08, 95% CI [0.70, 1.53], BF_10_ ≈ 7,549. Intrusions did not differ (*p* = .376). Recall declined more steeply from left to right under threat (group × column χ^2^(3) = 11.24, *p* = .011), with losses confined to the third and fourth letters of the top row (FDR-corrected *q* ≤ .003).

**Conclusions:** Anticipatory threat reduced the amount of information transferred from a brief display into a reportable form without increasing guessing. Intact first items and depleted later items along the habitual reading order indicate a reduced number of items read out of a rapidly decaying visual store, rather than a hemifield-specific bias. The single-trial design and the absence of a manipulation check limit mechanistic inference.

## 1. Introduction

Acute stress mobilises a cascade of neural and endocrine responses that unfold over different time scales. Within seconds, noradrenergic signalling reconfigures large-scale brain networks, enhancing vigilance and threat detection at the expense of executive control^1,2^; within minutes, glucocorticoids act on hippocampal and prefrontal circuits^3,4^. High levels of catecholamines rapidly impair the prefrontal functions that support working memory and attentional control^5^, and acute psychological stress reduces working memory–related activity in the dorsolateral prefrontal cortex^6^. At the behavioural level, stress before or during encoding generally impairs later memory, whereas stress after learning can enhance consolidation^7,8^, and the overall effect of psychosocial stressors on memory and related abilities is reliable but smaller than often assumed^9,10^.

Most of this work concerns working, episodic or long-term memory. Much less is known about the very first stage of visual storage. When a display is flashed briefly, observers have access to a high-capacity but rapidly decaying representation—iconic memory—from which only a limited number of items can be transferred into a durable, reportable form before it fades^11–13^. In whole-report tasks the number of items reported therefore reflects the efficiency of an attention-dependent read-out process operating under severe time pressure, and formal models describe performance in terms of a processing rate and a limited short-term storage capacity of roughly three to four items^14–17^. Because these parameters are sensitive to changes in attentional efficiency^18^, whole report is a promising probe of how anxiety constrains early information intake.

Anxiety is a natural candidate for disrupting this read-out. Attentional control theory proposes that anxiety reduces the efficiency of goal-directed attention by biasing resources towards threat^19^, and arousal has long been thought to narrow the range of cues that are utilised^20^ or to bias competition in favour of the most salient inputs^21^. Meta-analytic evidence links anxiety to reduced working memory capacity^22^, and induced anxiety selectively disrupts visuospatial working memory^23^. Anxious individuals also show threat-related attentional biases and weakened prefrontal control over distraction^24–26^. At the same time, emotion can enhance early perception^27^, so the net effect of anxiety on the first stages of visual processing cannot be assumed.

The threat-of-shock paradigm, in which participants anticipate an unpredictable aversive stimulus, reliably induces anticipatory anxiety and is a well-validated translational model of sustained fear^28–31^. Under threat, performance tends to suffer most when tasks place modest demands on attention, leaving resources free to be captured by the threat^32,33^—a profile that matches a single, brief whole-report trial.

Here we asked whether anticipatory threat reduces the number of items people can report from a 400-ms letter display. Beyond total recall, we examined *where* in the display losses occurred. If threat reduces read-out capacity, items reported late in the habitual left-to-right, top-to-bottom scanning order should be lost first. If threat instead biases spatial attention towards one hemifield, losses should concentrate on one side of the display regardless of scanning order. We separated correct recall from intrusion errors and used analyses appropriate to bounded count data and to letters nested within participants.

## 2. Materials and Methods

### 2.1. Participants

Eighty-eight healthy volunteers aged 18–25 years took part. Inclusion criteria were normal or corrected-to-normal vision, adequate literacy and no self-reported history of neurological, psychiatric or cognitive disorder. Participants were allocated to a control group (*n* = 44) or a threat-of-shock group (*n* = 44) by simple randomisation. With 44 participants per group, the study had 80% power (two-tailed α = .05) to detect a between-group effect of *d* ≥ 0.60. The study was approved by the Non-Interventional Research Ethics Committee of Üsküdar University (decision 2025-31, 27 March 2025) and conducted in accordance with the Declaration of Helsinki. All participants gave written informed consent and were debriefed after the session; no electrical stimulation was delivered.

### 2.2. Stimuli and task

The stimulus was a 7-s video presented on a single laboratory monitor. A black screen was shown for 5.5 s, followed by a 4 × 3 matrix of white upper-case letters on a black background for 400 ms (top row: K M R T; middle row: Z L A H; bottom row: C P S V), after which the screen returned to black (Figure 1). Letters were chosen to minimise chunking into words. The video was prepared in Adobe Premiere Pro with fixed resolution, font, size and luminance. Immediately afterwards, participants wrote down, on a blank sheet and without a time limit, every letter they could recall in any order (whole report with free written recall).

**Figure 1.**
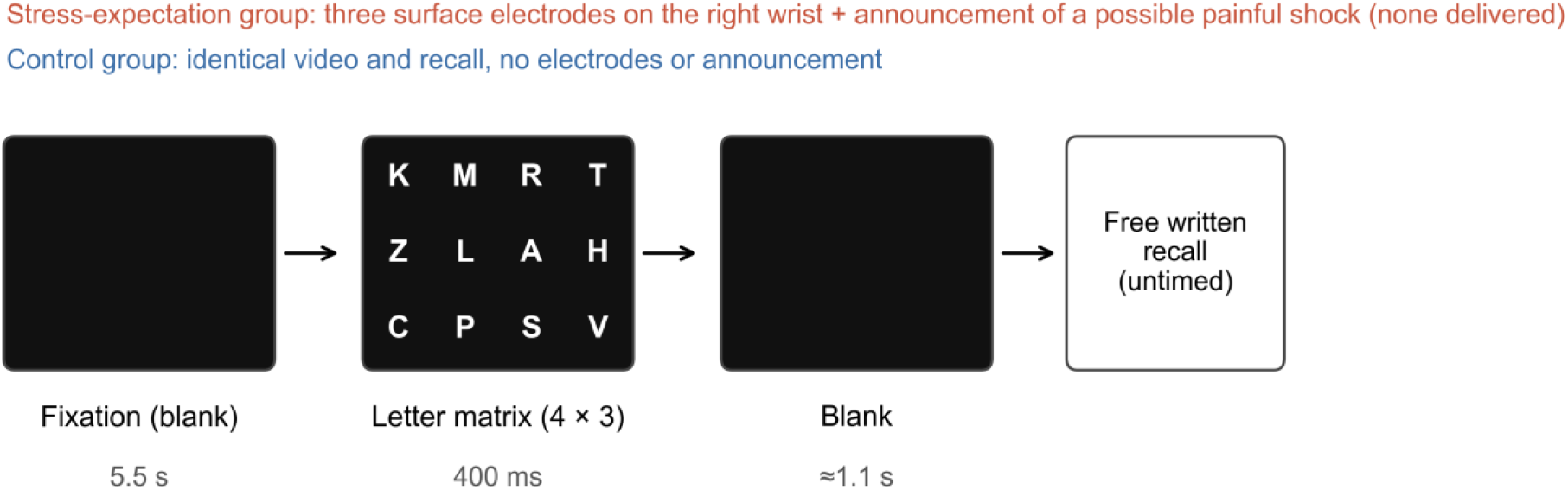
Trial structure. Both groups watched the same single-trial video and then performed untimed written free recall. In the threat-of-shock condition, three surface electrodes were attached to the right wrist before the video and participants were told that a painful electrical stimulus might be delivered during the task; no stimulus was delivered.

### 2.3. Threat-of-shock manipulation

In the threat condition, three surface electrodes were attached to the participant’s right wrist and the experimenter stated that a painful electrical stimulation might be applied at some point during the video. This instructed-threat procedure follows the logic of threat-of-shock paradigms, in which anticipation of an unpredictable aversive event is sufficient to induce anticipatory anxiety^28,31^. Participants were asked to watch the video and then to write down the letters as in the control condition. The control group received identical instructions without electrodes or any mention of stimulation. Testing took place individually in a quiet room of the Neuroscience Laboratory of Üsküdar University.

### 2.4. Scoring

Response sheets were transcribed verbatim. A letter was scored as correct if it was present in the matrix, irrespective of the position at which it was written, yielding a correct-recall score from 0 to 12. Letters that did not appear in the display were scored as intrusions; no participant reported a matrix letter more than once. Each of the twelve matrix positions was additionally coded as recalled (1) or not recalled (0) for every participant, allowing analysis of the spatial distribution of recall.

### 2.5. Statistical analysis

Normality was assessed with the Shapiro–Wilk test and homogeneity of variance with the median-centred Levene test. Because correct-recall scores deviated from normality, the primary comparison used Welch’s *t* test, which is robust to unequal variances^34^, and was confirmed with the Mann–Whitney *U* test (effect size: rank-biserial *r*). Hedges’ *g* and the mean difference are reported with 95% percentile bootstrap confidence intervals (10,000 resamples)^35^, and evidence for the alternative hypothesis was quantified with the default JZS Bayes factor (BF_10_)^36^. Because recall is a bounded count, a binomial generalised linear model (12 trials per participant; logit link) was also fitted, and the odds ratio (OR) for the threat condition is reported.

Spatial effects were analysed at the level of individual matrix positions (12 binary observations per participant) using generalised estimating equations with a logit link and an exchangeable working correlation structure clustered by participant^37^. Separate models tested group × column (four levels) and group × row (three levels) interactions with Wald χ^2^ tests; a further model entered column as a linear term to estimate the change in the left-to-right gradient under threat. Group differences for each letter were tested with Fisher’s exact tests, controlling the false discovery rate with the Benjamini– Hochberg procedure (*q*)^38^. As a sensitivity analysis, the main comparison was repeated after excluding the two letters (K and M) that were recalled by almost all participants. Analyses were run in Python 3.10 (SciPy, statsmodels, pingouin) with α = .05 (two-tailed).

## 3. Results

### 3.1. Total correct recall

Correct-recall scores were non-normally distributed in both groups (control: *W* = 0.875, *p* < .001; threat: *W* = 0.893, *p* < .001), whereas variances were homogeneous (Levene *F* = 0.11, *p* = .741). Participants under threat of shock reported fewer correct letters than controls (Table 1; Figure 2): Welch *t*(83.6) = 5.11, *p* < .001, mean difference = 1.30 letters, bootstrap 95% CI [0.82, 1.80]; Hedges’ *g* = 1.08 [0.70, 1.53]. The non-parametric test led to the same conclusion (*U* = 1508, *p* < .001, rank-biserial *r* = 0.56), and the Bayes factor indicated extreme evidence for a group difference (BF_10_ ≈ 7,549). In the binomial model, the odds of reporting any given letter were 38% lower under threat (OR = 0.62, 95% CI [0.48, 0.80], *p* < .001; no overdispersion, Pearson χ^2^/*df* = 0.53).

**Table 1.** Correct Recall and Intrusion Errors by Group.

| **Measure** | **Control (n = 44)** | **Threat of shock (n = 44)** | **Test** | **p** |
|---|---|---|---|---|
| Correct letters, M (SD) | 4.70 (1.29) | 3.41 (1.09) | Welch t(83.6) = 5.11 | < .001 |
| Correct letters, Mdn [IQR] | 4 [4–5] | 3 [3–4] | U = 1508 | < .001 |
| Range | 2–8 | 2–6 |  |  |
| Intrusions, M (total) | 0.34 (15) | 0.20 (9) | U = 1044 | .376 |
| Letters written, M | 5.05 | 3.61 |  |  |

*Note.* Correct recall = number of displayed letters reported (0–12). Effect sizes for correct recall: Hedges' g = 1.08, 95% bootstrap CI [0.70, 1.53]; rank-biserial r = 0.56; BF<sub>10</sub> ≈ 7,549. IQR = interquartile range.

**Figure 2.**
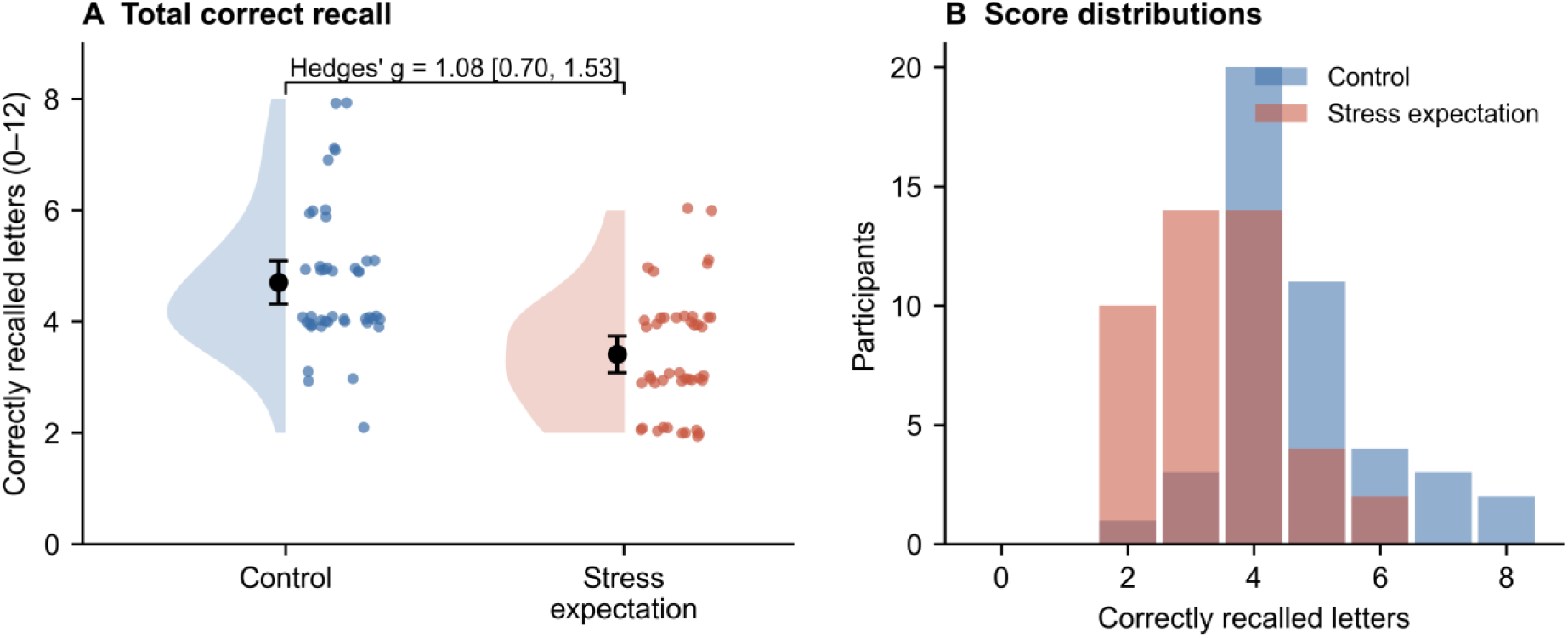
Correct recall by group. (A) Individual scores (points; vertical jitter added for visibility), distribution (half-violin) and group mean with 95% confidence interval (black). (B) Frequency distribution of scores.

### 3.2. Intrusion errors

Intrusions were rare and did not differ between groups (15 in total in the control group vs. 9 in the threat group; *U* = 1044, *p* = .376, rank-biserial *r* = 0.08). The threat-related deficit therefore reflects fewer correctly reported items rather than a more liberal or noisier response strategy.

### 3.3. Spatial distribution of recall

Both groups showed a pronounced top-to-bottom gradient, with the first letters of the top row reported by almost every participant (K: 43/44 vs. 43/44; M: 42/44 vs. 41/44) and bottom-row letters rarely reported (Figure 3). The threat-related loss, however, was not uniform. The left-to-right decline in recall was steeper under threat (group × column: Wald χ^2^(3) = 11.24, *p* = .011; linear group × column OR = 0.74, 95% CI [0.61, 0.89], *p* = .001), and the difference was largest in the rightmost column (interaction OR relative to column 1 = 0.35, *p* < .001; Figure 4A). The group × row interaction was not significant (χ^2^(2) = 4.41, *p* = .110; Figure 4B).

**Figure 3.**
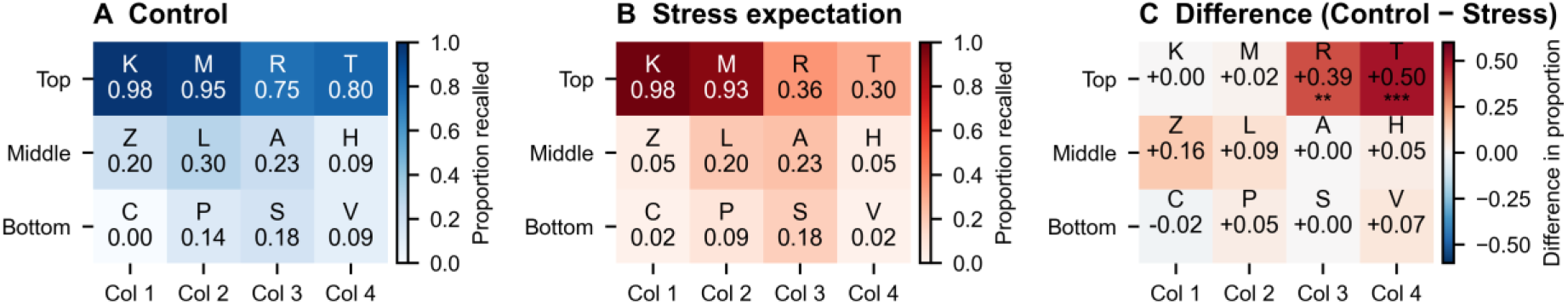
Spatial distribution of recall. Proportion of participants reporting each letter in the control (A) and threat-of-shock (B) groups, and the difference between groups (C). Asterisks in (C) mark letters whose group difference survived Benjamini–Hochberg correction across the 12 positions (Fisher’s exact tests; *q* < .01, q *<* .*001)*.

**Figure 4.**
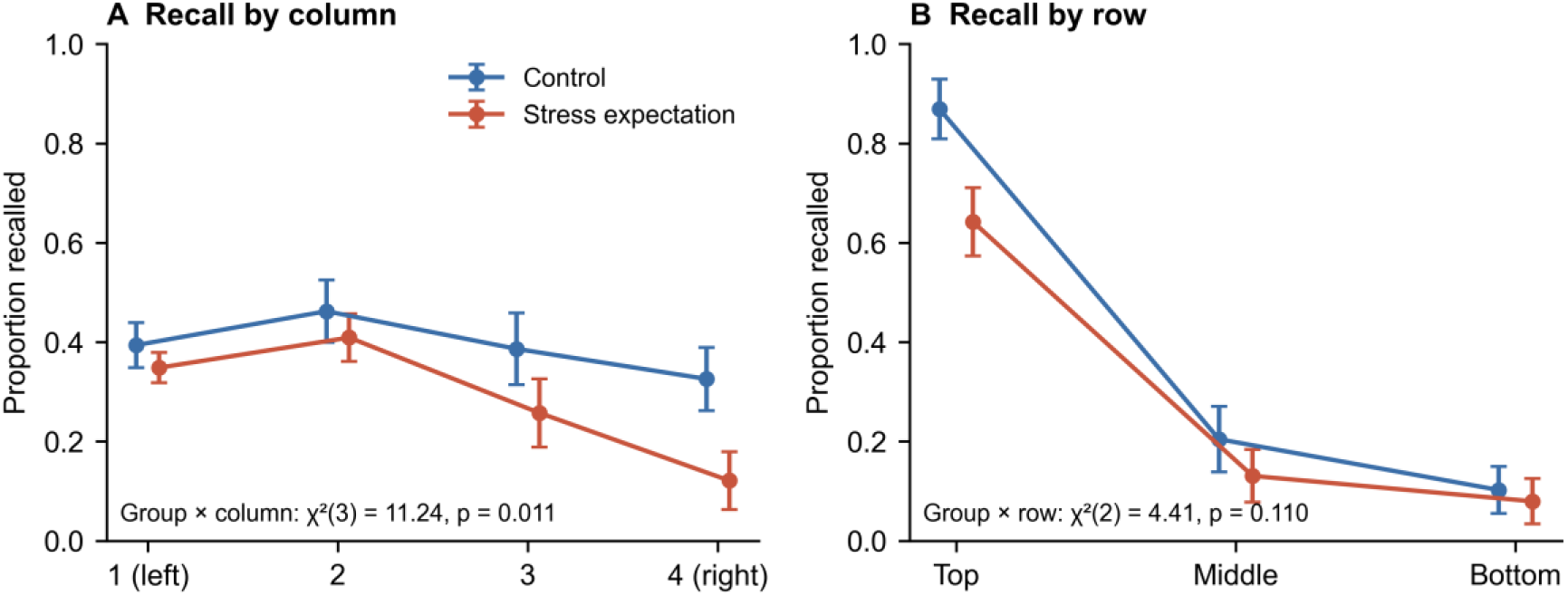
Recall as a function of display column (A) and row (B). Points show the mean of participant-level proportions with 95% confidence intervals. Statistics are Wald tests of the group × position interaction from generalised estimating equations (logit link, exchangeable correlation within participant).

At the level of individual letters, only the third and fourth items of the top row differed after FDR correction: R (33/44 vs. 16/44, *q* = .003) and T (35/44 vs. 13/44, *q* < .001). Z, the first letter of the middle row, showed a nominal difference (9 vs. 2, uncorrected *p* = .049) that did not survive correction (Table 2). Excluding K and M, which were at ceiling, left the group difference unchanged (2.77 vs. 1.50 letters; *U* = 1506, *p* < .001).

**Table 2.** Recall of Each Matrix Position by Group.

| **Letter (position)** | **Control, n (%)** | **Threat, n (%)** | **p** | **q (FDR)** |
|---|---|---|---|---|
| K (row 1, col 1) | 43 (98%) | 43 (98%) | 1.000 | 1.000 |
| M (row 1, col 2) | 42 (95%) | 41 (93%) | 1.000 | 1.000 |
| R (row 1, col 3) | 33 (75%) | 16 (36%) | < .001 | .003 |
| T (row 1, col 4) | 35 (80%) | 13 (30%) | < .001 | < .001 |
| Z (row 2, col 1) | 9 (20%) | 2 (5%) | .049 | .197 |
| L (row 2, col 2) | 13 (30%) | 9 (20%) | .461 | 1.000 |
| A (row 2, col 3) | 10 (23%) | 10 (23%) | 1.000 | 1.000 |
| H (row 2, col 4) | 4 (9%) | 2 (5%) | .676 | 1.000 |
| C (row 3, col 1) | 0 (0%) | 1 (2%) | 1.000 | 1.000 |
| P (row 3, col 2) | 6 (14%) | 4 (9%) | .739 | 1.000 |
| S (row 3, col 3) | 8 (18%) | 8 (18%) | 1.000 | 1.000 |
| V (row 3, col 4) | 4 (9%) | 1 (2%) | .360 | 1.000 |

*Note*. Fisher's exact tests; q = Benjamini–Hochberg-adjusted p across the 12 positions.

## 4. Discussion

Anticipating a possible electric shock reduced the number of letters participants could report from a single 400-ms display by about 1.3 letters, a large effect (*g* ≈ 1.08) that was consistent across parametric, non-parametric, Bayesian and count-based analyses. Intrusions did not increase, indicating that the deficit reflects less information being transferred into a reportable form rather than greater guessing or response bias. Relative to the control mean of about 4.7 items—close to the three-to-five-item capacity estimates of visual short-term memory^16,17^—threat reduced report to roughly 3.4 items.

The spatial pattern constrains how the deficit should be interpreted. In both groups, report followed the habitual reading order: the first letters of the top row were reported almost universally and performance fell steeply thereafter. Threat did not affect these first items; it depleted the items that come next in a left-to-right, top-to-bottom scan—most clearly the third and fourth letters of the top row. Although this produced a right-sided loss, the deficit did not extend to the right-hand letters of the lower rows. The more parsimonious account is therefore a reduction in the number of items read out of the decaying visual store before it fades^11,13^. In the framework of the theory of visual attention, such a pattern would follow from a lower processing rate or a shorter effective exposure, rather than from a change in attentional weighting across space^14,15,18^. This interpretation is consistent with attentional control theory, which predicts reduced processing efficiency under anxiety^19^, with the observation that anxiety selectively impairs visuospatial storage^23^, and with classic and contemporary accounts of arousal-related narrowing of information intake^20,21^.

A hemifield-specific account cannot be excluded entirely. Healthy observers show a slight leftward attentional bias (pseudoneglect)^39^, and anxiety-related changes in arousal systems could in principle exaggerate such asymmetries. Distinguishing between the two accounts requires designs that dissociate scanning order from visual field—for example, displays read in the opposite direction, partial-report cueing^11,12^ or eye tracking.

The timing of the manipulation is also informative. Threat was induced seconds to minutes before the display, a window dominated by rapid noradrenergic and anticipatory anxiety responses that shift network balance towards salience and vigilance at the expense of executive control^1,2,5^, rather than by peak glucocorticoid effects^3,4^. Threat of shock reliably engages sustained anxiety circuitry^29,30^ and shifts processing towards threat monitoring, particularly when task load is low^32,33^. A single brief display fits this profile, and the results suggest that anticipatory anxiety constrains information intake at the earliest stage at which visual information becomes available for report. These findings complement evidence that anxiety biases attention towards threat and weakens prefrontal control over distractors^24–26^, and indicate that such costs are not confined to working memory maintenance^6,22^.

### 4.1. Limitations

Several limitations qualify these conclusions. First, no manipulation check was obtained: neither subjective anxiety ratings nor physiological indices (e.g., skin conductance, heart rate, startle or salivary cortisol) were recorded, so the magnitude of the induced state is unknown and individual differences in threat reactivity could not be modelled^31^. Second, each participant completed a single trial; whole-report estimates from one trial are noisy, and the design did not allow us to separate iconic decay from later working-memory processes or to estimate processing-rate and capacity parameters^14^. Third, the experimenter was aware of group allocation, and the electrode procedure itself may have introduced distraction independent of anxiety; an electrodes-without-threat control would isolate the threat component. Fourth, the same letter matrix was used for all participants, so letter identity and position are confounded. Finally, the sample consisted of young adults, and sex and other demographic variables were not analysed.

### 4.2. Conclusions and future directions

Anticipatory threat of shock reduces the amount of information that can be reported from a briefly presented visual display, selectively eliminating items that would be processed late in the habitual scanning order. Future studies should combine multi-trial whole- and partial-report procedures with manipulation checks, physiological arousal measures and eye tracking, and should test whether the effect generalises to other anxiety inductions and to settings in which brief visual information must be acquired under pressure.

## Declarations

### Ethics statement

Approved by the Non-Interventional Research Ethics Committee of Üsküdar University (decision 2025-31, 27 March 2025). Written informed consent was obtained from all participants, who were debriefed after the session.

### Data availability

Anonymised participant-level data and the analysis script will be deposited in a public repository (Open Science Framework) upon acceptance and are available from the corresponding author on request.

### Author contributions (CRediT)

Barış Akar: Conceptualization, Methodology, Investigation, Formal analysis, Visualization, Writing – original draft. Merve Aydın Karabaş: Investigation, Writing – review & editing. Tuğba Altıntaş: Methodology, Supervision, Writing – review & editing.

### Use of generative AI

A generative AI tool was used to assist with re-analysis, figure preparation and language editing. The authors verified all analyses and text and take full responsibility for the manuscript.

